# Novel Pentafluorosulfanyl-containing Triclocarban Analogs selectively kill Gram-positive bacteria

**DOI:** 10.1101/2024.01.04.574235

**Authors:** Ali Pormohammad, Melika Moradi, Josefien W. Hommes, Eugènia Pujol, Lieve Naesens, Santiago Vázquez, Bas G. J. Surewaard, Mohammad Zarei, Manuel Vazquez-Carrera, Raymond J. Turner

## Abstract

The antibacterial and antibiofilm efficacy of our novel pentafluorosulfanyl-containing triclocarban analogs was explored against seven different Gram-positive and Gram-negative indicator strains. After initial screening, they had bactericidal and bacteriostatic activity against Gram-positive bacteria, especially *Staphylococcus aureus* and *MRSA* (methicillin–resistant staphylococcus aureus) in a very low concentration. Our results were compared with the most common antibiotic being used (Ciprofloxacin and Gentamycin); the novel components had significantly better antibacterial and antibiofilm activity in lower concentrations in comparison to the antibiotics. For instance, EBP-**59** minimum inhibitory concentration was < 0.0003 mM, while ciprofloxacin 0.08 mM. Further antibacterial activity of novel components was surveyed against 10 clinical antibiotic resistance *MRSA* isolates. Again, novel components had significantly better antibacterial and antibiofilm activity in comparison with antibiotics. Mechanistic studies have revealed that none of these novel compounds exhibit any effect on the reduced thiol, disrupting iron sulfur clusters, or hydrogen peroxide pathways. Instead, their impact is attributed to the disruption of the Gram-positive bacterial cell membrane. Toxicity and safety testing on tissue cell culture showed promising results for the safety of components to the host.

## Introduction

Bacterial resistance to antimicrobials has become a pressing concern in healthcare, demanding the development of effective formulations targeting multi-resistant pathogens (1). To address this challenge, researchers have turned to drug repurposing, a strategy that explores novel uses for authorized or experimental pharmaceuticals beyond their initial applications, offering advantages such as reduced costs and shorter development timelines (2). In this context, the exploration of multitarget molecules has emerged as a promising approach to counteract pathogens through diverse mechanisms. Diarylureas, such as regorafenib, sorafenib, linifanib, ripretinib, and tivozanib, have long been recognized as anticancer agents (3). One intriguing avenue involves the repurposing of diarylureas with anticancer properties for novel indications, such as antimicrobial, anti-inflammatory, and antiviral applications (4).

Trifluoromethyl groups, serving as bio-isosteric replacements for chlorine atoms, have found widespread use in medicinal chemistry. Hence, it is unsurprising that specific N,N′-diarylureas that comprise a trifluoromethyl moiety have demonstrated encouraging antibacterial properties. (5). An example is cloflucarban (TFC, 3-trifluoromethyl-4,4′-dichlorocarbanilide), a trifluoromethyl-substituted diarylurea which shares a similar spectrum of activity and pharmacokinetic profile with triclocarban (TCC). In recent times, a number of diarylureas analogs of TCC have been identified as having antibacterial and antifungal properties. The presence of pentafluorosulfanyl and trifluoromethylcoumarine groups characterizes these analogs. Given the risks associated with TCC, the need for alternative antimicrobial agents has become crucial (3).

In the recent years, a novel bio-isosteric trifluoromethyl unit, the pentafluorosulfanyl group (SF_5_), has emerged in medicinal chemistry, finding applications in agriculture and material chemistry (6–8). Regarded as a “super-trifluoromethyl group,” the SF_5_-group possesses several advantageous properties over its isostere trifluoromethyl group, the compound exhibits a tetragonal bipyramidal morphology and possesses a greater electronegativity value of 3.65, in comparison to trifluoromethyl’s value of 3.36. Additionally, it displays higher lipophilicity and notable steric volume, which is a bit smaller compared to *tert*-butyl but more than trifluoromethyl. The compound’s hydrolytic and chemical stability has also been approved (9). These unique characteristics have led to a significant rise in the utilization of SF_5_ in medicinal chemistry over the past decade, making it an exceptionally appealing substituent for medicinal applications. SF_5_-containing building blocks have garnered attention among medicinal chemists owing to their stated ability to decelerate metabolic rates and their eco-friendliness vis-à-vis lack of chlorine atoms, despite their elevated cost relative to analogous CF_3_ compounds. Building upon the growing utilization of SF_5_ in medicinal chemistry and its favorable environmental profile, the objective of this study is to incorporate this innovative group onto the N,N′-diarylurea scaffold to explore for novel antimicrobial agents (10).

This present study aims to explore the synthesis and antimicrobial and antibiofilm activity of novel diphenylurea agents, particularly inspired by TCC but bearing different aryl moieties. Mechanistic studies, such as cell wall or hydrogen peroxide assays, are conducted to elucidate the compounds’ modes of action. Cytotoxicity assays using cell lines will ensure their safety as a unique class of antimicrobials.

## Result and Discussion

During the recent years, overuse, and misuse of antibiotics against bacterial infections has brought up an issue regarding prevalence of multi-resistant bacteria to conventional antibiotics. This has led that development of novel antimicrobial agents and their delivery mechanism has turned into a global concern (11). Given that diarylureas analogs of TCC have been recognized for their antibacterial and antifungal properties (3), our study investigates the antibacterial efficacy of our novel pentafluorosulfanyl-containing triclocarban analogs.

### Chemistry

The eighteen *N,N’*-diarylureas evaluated in this work were synthesized following a simple and straightforward procedure consisting of the coupling of phenyl isocyanates with the corresponding anilines, as previously described by some of us reported (5, 12, 13) (**Fig 1**). In turn, the intermediate phenyl isocyanates were either commercially available or synthesized *in situ* from the reaction of the precursor anilines with triphosgene. The analytical data of the compounds fully agreed with the data previously reported (5, 12, 13).

**Fig 1.**
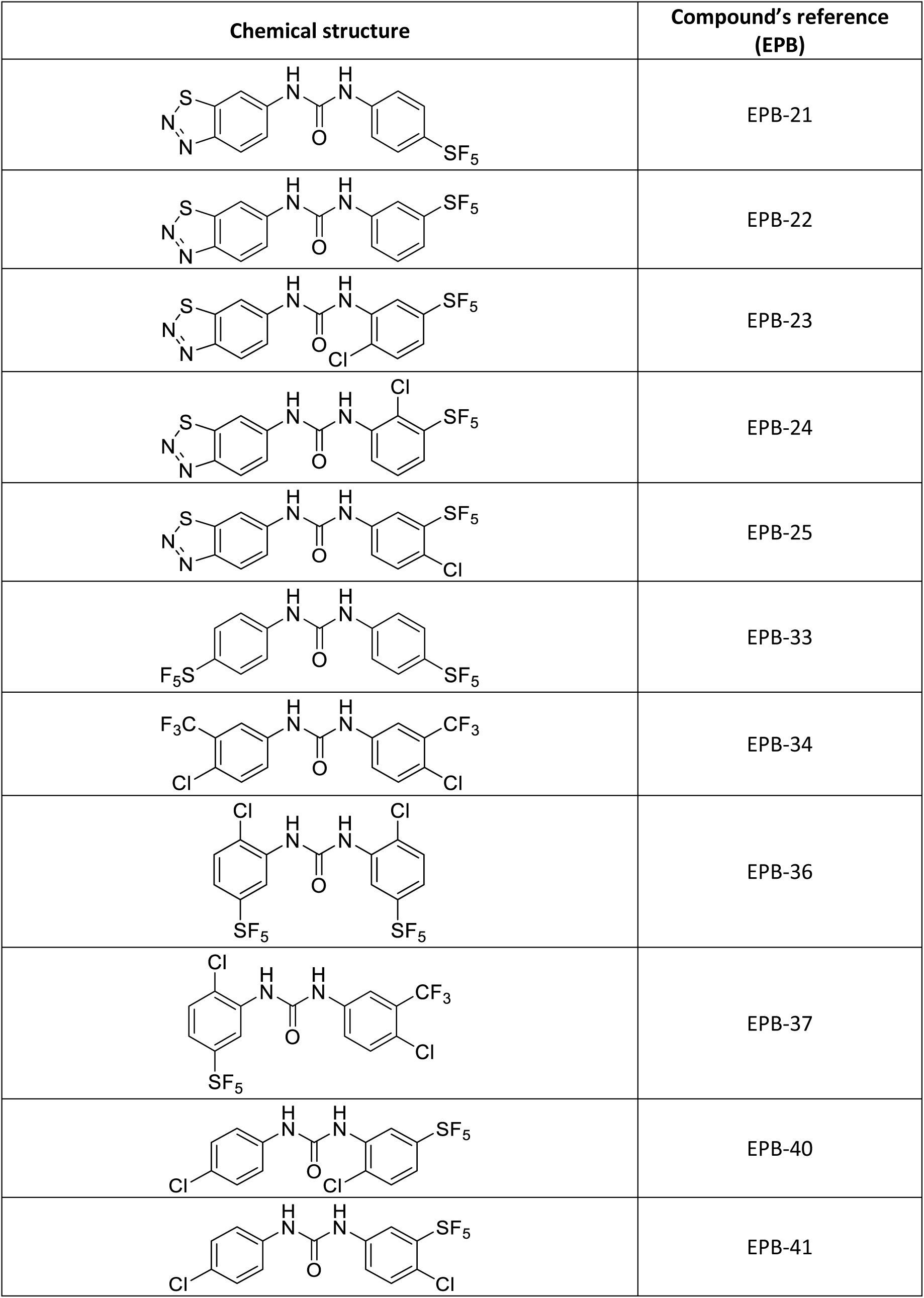
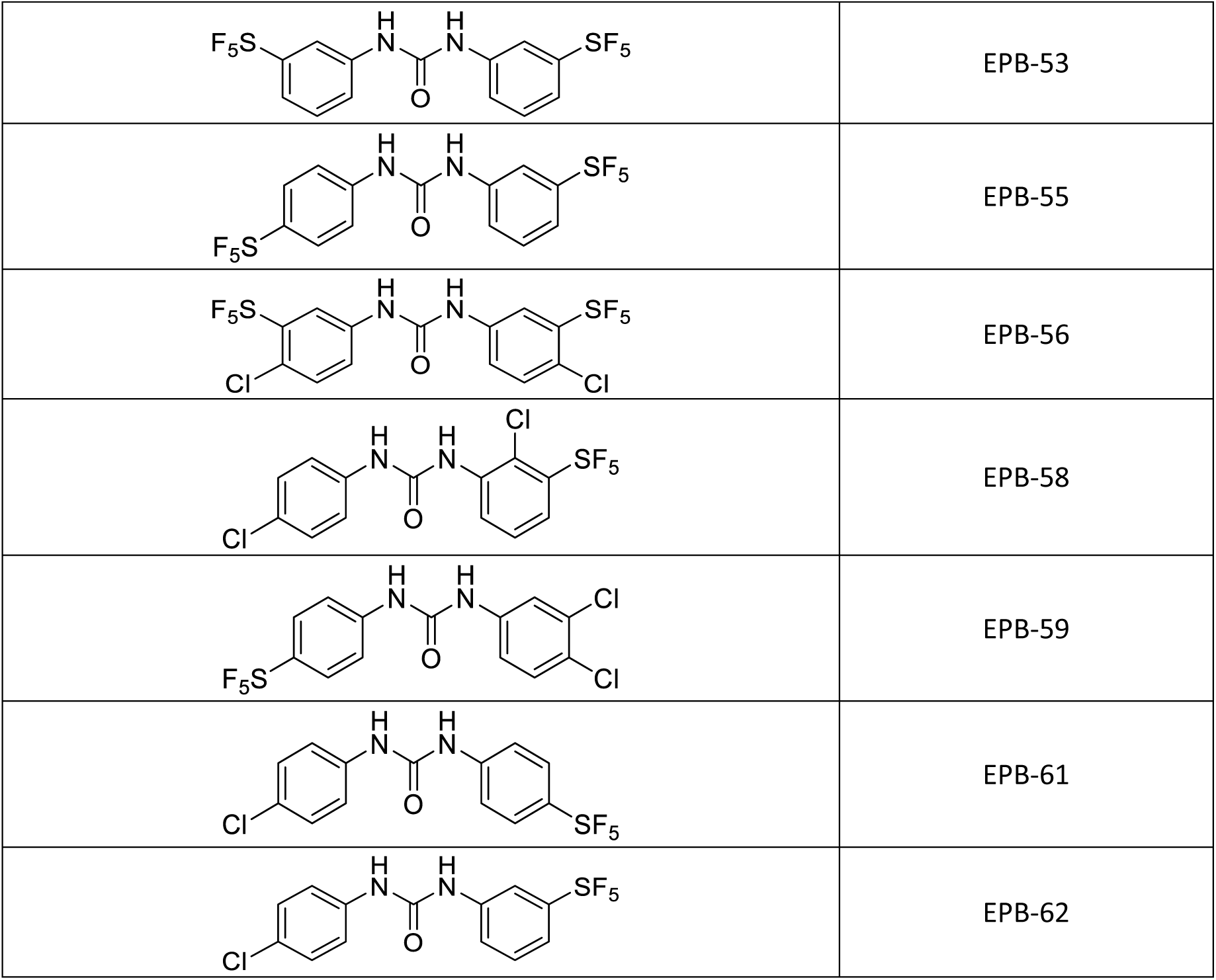
Compounds EPB-21, EPB-22, EPB-23, EPB-24, EPB-25, EPB-36 (12), compounds EPB-33, PB-34, EPB-37, EPB-40, EPB-41, EPB-55, EPB-56, EPB-58, EPB-59, EPB-61, EPB-62 (5, 12) and EPB-53 have been published in our previous works (5, 28).

### Antibacterial activity

We explored the antibacterial activity of compounds against several bacterial pathogens, obtaining measurements of minimum inhibitory concentration (MIC), and minimum bactericidal concentration (MBC), (**Table 1**). None of the diphenylurea compounds were effective against any of the Gram-negative bacteria evaluated in this study. The only exception was the effect of compound **59** on *P. mirabilis* with MIC and MBC of 0.002 mM. However, regarding the main outcome measure of this study, three compounds **59**, **61**, and **62** were effective against *S. aureus, S. epidermidis* and *MRSA* spp. Exact dosages in respect to each compound and bacteria is illustrated in **Table1**. Compound **59** showed the best efficacy against *S. aureus*, *MRSA*, and *S. epidermidis*. Compound **61** also was effective against these bacteria with relatively higher doses. Compound **62** was also effective against these Gram-positive bacteria, however, it had the highest doses comparatively (**Fig 2**).

**Fig 2.**
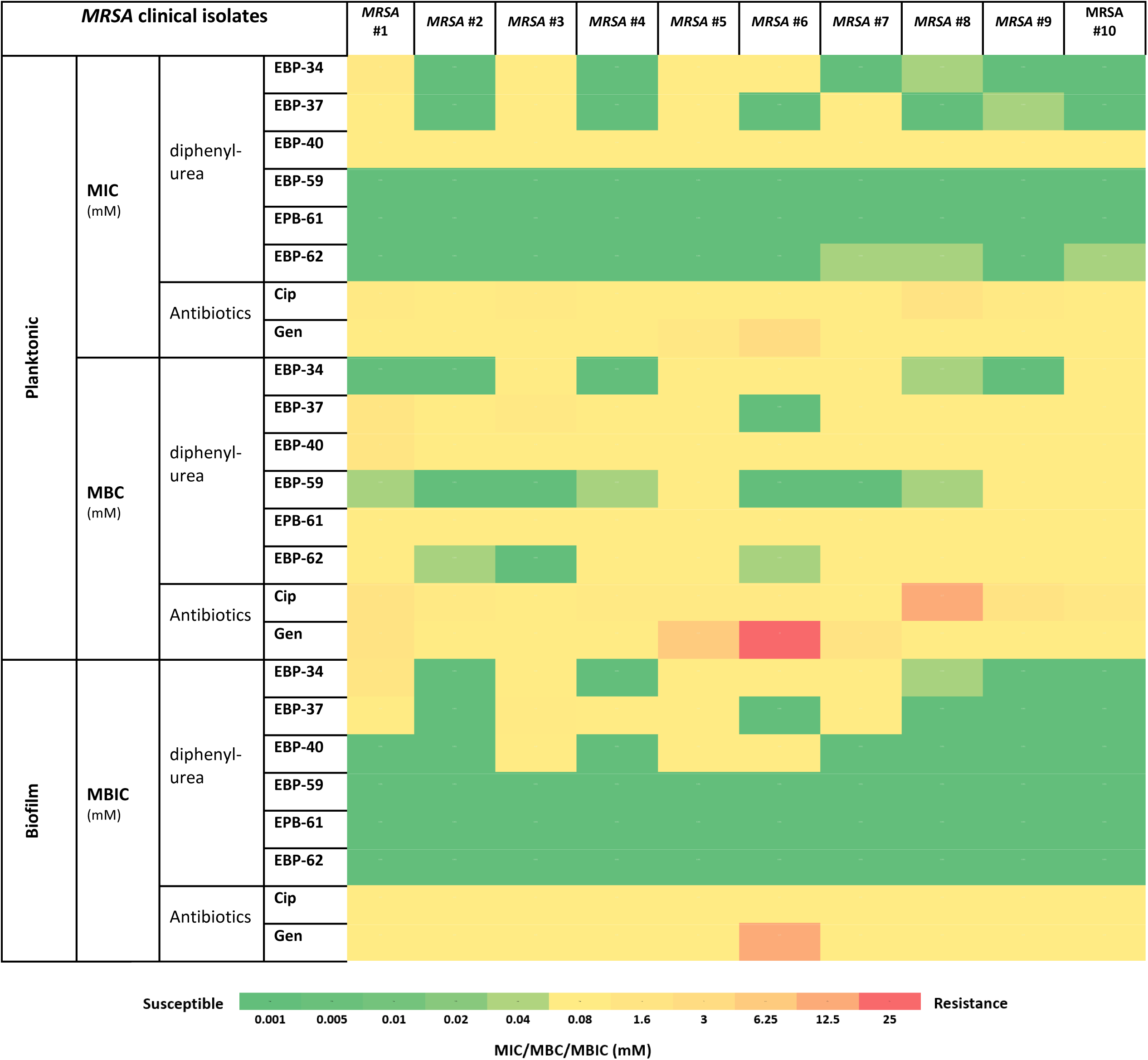
Minimum inhibitory concentration (MIC), Minimum bactericidal concentration (MBC), and Minimum biofilm inhibition concentration (MBIC) of diphenyl-urea antibiotics against *MRSA* clinical isolates.

**Table 1.**
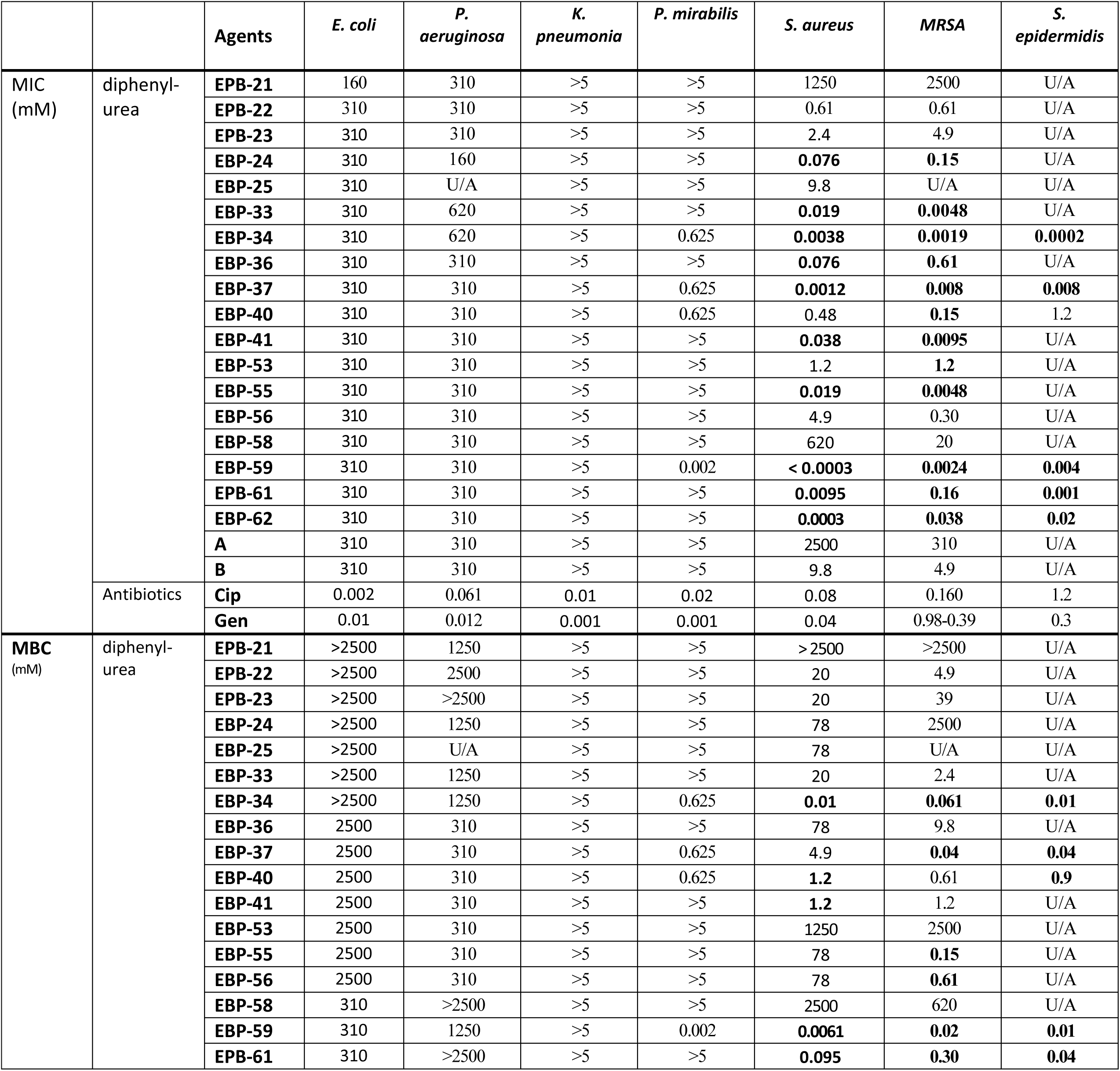
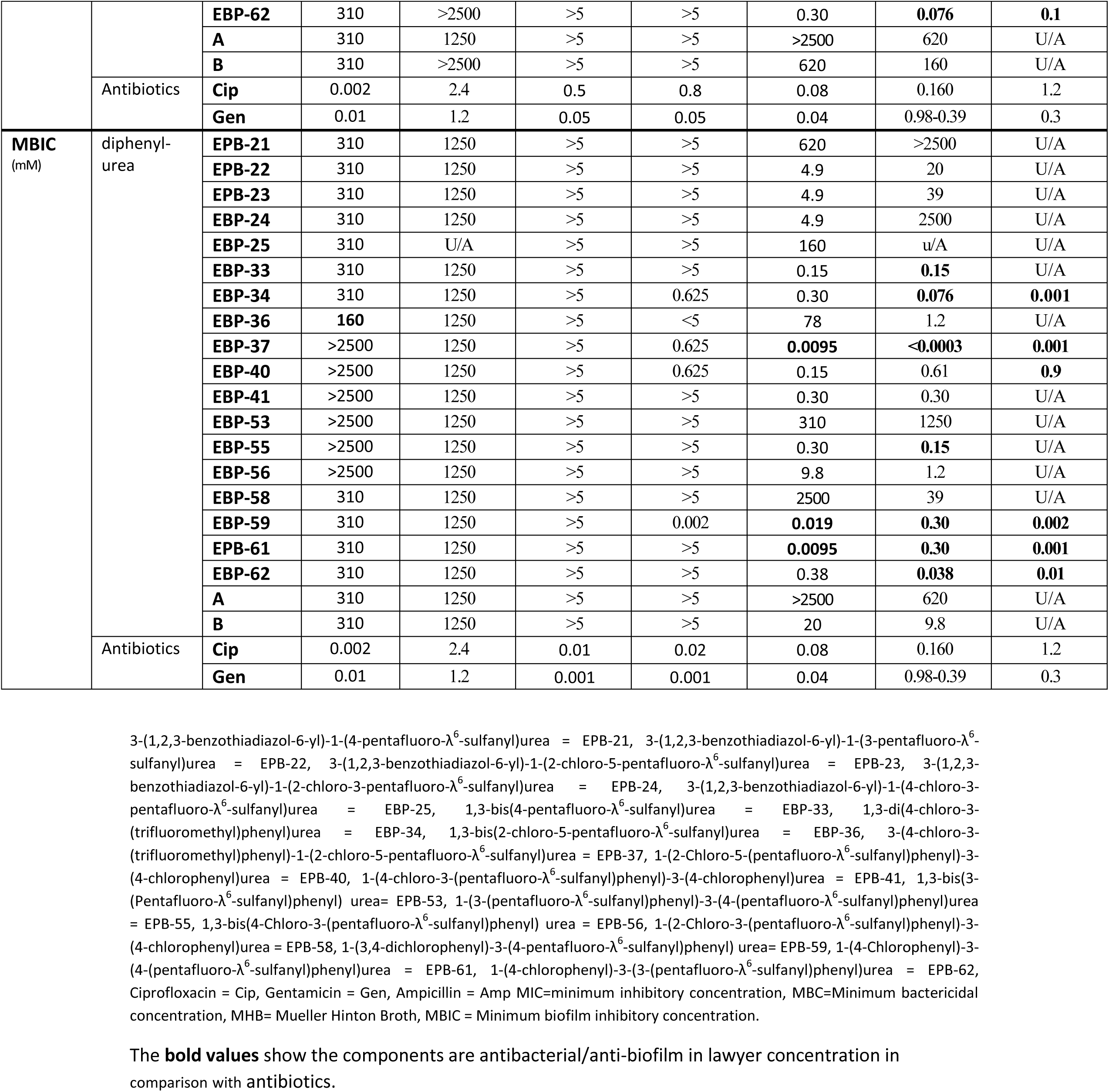
Minimum inhibitory concentration (MIC), Minimum bactericidal concentration (MBC) and Minimum biofilm inhibitory concentration (MBIC) of diphenyl-urea antibiotics for *P. aeruginosa*, *S. aureus* and *E. coli*.

Given the efficacy against the reference strain of MSRA, we tested further ten clinical isolates of *MRSA* evaluated with compounds **59**, **61** and **62**. The results were extremely promising showing similarly effective doses of MIC and MBC against the clinical isolates (**Table 2**).

**Table 2.** Minimum inhibitory concentration (MIC), Minimum bactericidal concentration (MBC) and Minimum biofilm inhibitory concentration (MBIC) of diphenyl-urea antibiotics against *MRSA* clinical isolates.

| MRSA clinical isolates |  |  | *MRSA*<br>#1 | *MRSA*<br>#2 | *MRSA*<br>#3 | *MRSA*<br>#4 | *MRSA*<br>#5 | *MRSA*<br>#6 | *MRSA*<br>#7 | *MRSA*<br>#8 | *MRSA*<br>#9 | *MRSA*<br>#10 |
| --- | --- | --- | --- | --- | --- | --- | --- | --- | --- | --- | --- | --- |
| **MIC** (mM) | diphenyl-urea | **EBP-34** | 0.6 | < 0.002 | 0.08 | < 0.002 | 0.01 | 0.01 | < 0.002 | 0.005 | < 0.002 | < 0.002 |
|  |  | **EBP-37** | 0.01 | < 0.002 | 0.08 | < 0.002 | 0.01 | < 0.002 | 0.01 | < 0.002 | 0.005 | < 0.002 |
|  |  | **EBP-40** | 0.02 | 0.02 | 0.08 | 0.04 | 0.02 | 0.04 | 0.04 | 0.04 | 0.02 | 0.04 |
|  |  | **EBP-59** | < 0.002 | < 0.002 | < 0.002 | < 0.002 | < 0.002 | < 0.002 | < 0.002 | < 0.002 | < 0.002 | < 0.002 |
|  |  | **EPB-61** | < 0.002 | < 0.002 | < 0.002 | < 0.002 | < 0.002 | < 0.002 | < 0.002 | < 0.002 | < 0.002 | < 0.002 |
|  |  | **EBP-62** | < 0.002 | < 0.002 | < 0.002 | < 0.002 | < 0.002 | < 0.002 | 0.005 | 0.005 | < 0.002 | 0.005 |
|  | Antibiotics | **Cip** | 0.4 | 0.05 | 0.4 | 0.05 | 0.2 | 0.2 | 0.2 | 1.6 | 0.4 | 0.05 |
|  |  | **Gen** | 0.1 | 0.05 | 0.05 | 0.05 | 0.8 | 3 | 0.05 | 0.05 | 0.05 | 0.05 |
| **MBC** (mM) | diphenyl-urea | **EBP-34** | < 0.002 | < 0.002 | 0.08 | < 0.002 | 0.01 | 0.01 | 0.01 | 0.005 | < 0.002 | 0.01 |
|  |  | **EBP-37** | 1.2 | 0.01 | 0.6 | 0.01 | 0.01 | < 0.002 | 0.04 | 0.01 | 0.01 | 0.01 |
|  |  | **EBP-40** | 1.2 | 0.04 | 0.16 | 0.16 | 0.04 | 0.16 | 0.08 | 0.08 | 0.08 | 0.01 |
|  |  | **EBP-59** | 0.005 | < 0.002 | < 0.002 | 0.005 | 0.01 | < 0.002 | < 0.002 | 0.005 | 0.02 | 0.04 |
|  |  | **EPB-61** | 0.04 | 0.01 | 0.01 | 0.02 | 0.04 | 0.02 | 0.02 | 0.04 | 0.04 | 0.02 |
|  |  | **EBP-62** | 0.08 | 0.005 | < 0.002 | 0.02 | 0.02 | 0.005 | 0.01 | 0.08 | 0.04 | 0.08 |
|  | Antibiotics | **Cip** | 1.6 | 0.4 | 0.05 | 0.4 | 0.4 | 0.4 | 0.05 | 12.5 | 1.6 | 0.8 |
|  |  | **Gen** | 1.6 | 0.2 | 0.05 | 0.05 | 6.25 | 25 | 1.6 | 0.05 | 0.05 | 0.05 |
| **MBIC** (mM) | diphenyl-urea | **EBP-34** | 1.2 | < 0.002 | 0.08 | < 0.002 | 0.01 | 0.01 | 0.02 | 0.005 | < 0.002 | < 0.002 |
|  |  | **EBP-37** | 0.02 | < 0.002 | 0.3 | 0.01 | 0.01 | < 0.002 | 0.01 | < 0.002 | < 0.002 | < 0.002 |
|  |  | **EBP-40** | < 0.002 | < 0.002 | 0.08 | < 0.002 | 0.02 | 0.04 | < 0.002 | < 0.002 | < 0.002 | < 0.002 |
|  |  | **EBP-59** | < 0.002 | < 0.002 | < 0.002 | < 0.002 | < 0.002 | < 0.002 | < 0.002 | < 0.002 | < 0.002 | < 0.002 |
|  |  | **EPB-61** | < 0.002 | < 0.002 | < 0.002 | < 0.002 | < 0.002 | < 0.002 | < 0.002 | < 0.002 | < 0.002 | < 0.002 |
|  |  | **EBP-62** | < 0.002 | < 0.002 | < 0.002 | < 0.002 | < 0.002 | < 0.002 | < 0.002 | < 0.002 | < 0.002 | < 0.002 |
|  | Antibiotics | **Cip** | 0.05 | 0.05 | 0.2 | 0.05 | 0.05 | 0.2 | 0.05 | 0.05 | 0.05 | 0.05 |
|  |  | **Gen** | 0.05 | 0.05 | 0.05 | 0.05 | 0.1 | 12.5 | 0.05 | 0.05 | 0.05 | 0.05 |

### Antibiofilm activity

The antibiofilm activity of antimicrobials can vary significantly compared to their effectiveness against planktonic forms of bacteria due to differences in physiology and structure of the biofilm (14). In fact, certain reports have demonstrated that biofilms require concentrations of antibiotics up to 100 times higher than those needed to eliminate planktonic bacteria (15–17). To assess the antibiofilm potential of the novel compounds, their Minimum Biofilm Inhibitory Concentration (MBIC) were determined **Table 1**. Similar to planktonic antibacterial activity, compounds **59**, **61** and **62** were effective against biofilm form of Gram-positive bacteria. Compounds **59** and **61** had the best antibiofilm activity in comparison to other compounds against *S. epidermidis, MRSA* and *S. aureus*. Compound **62** also showed relatively better results against these bacteria. When evaluating these compounds against ten clinical isolates of *MRSA*, all these isolates showed had their biofilms inhibited by these compounds (**Table 2**).

### Antimicrobial mechanism of action

A wide variety of mechanisms of antimicrobial action are reported for antimicrobial agents. Here in this study the most common mechanisms such as oxidation of the redox buffer reflected in reduced thiol content (19), related oxidative stress as seen by increase in reactive oxygen species (20) and breakdown of redox enzyme [Fe-S] clusters (21), and cell membrane dysfunction (11) were explored to have a general view of the various diphenylurea compounds mechanism of action.

A well-established method to evaluate bacteria viability, propidium iodide (PI) staining was performed to assess the membrane permeability. Compounds **59** and **61** showed no effect on the cell membrane of Gram-negative bacteria, *P. aeruginosa*. However, both compounds were effective at cell membrane disruption of *MRSA* strains. Proving the effective bactericidal of such compounds on the Gram-positive bacteria’s cell membrane (**Fig 3**). Further evaluations including reduced thiol, iron detection Ferine-S, and hydrogen peroxide levels were performed to explore other possible mechanisms of action of these compounds. These assays demonstrated that none of these novel compounds have any effect on these processes of toxicity.

**Fig 3.**
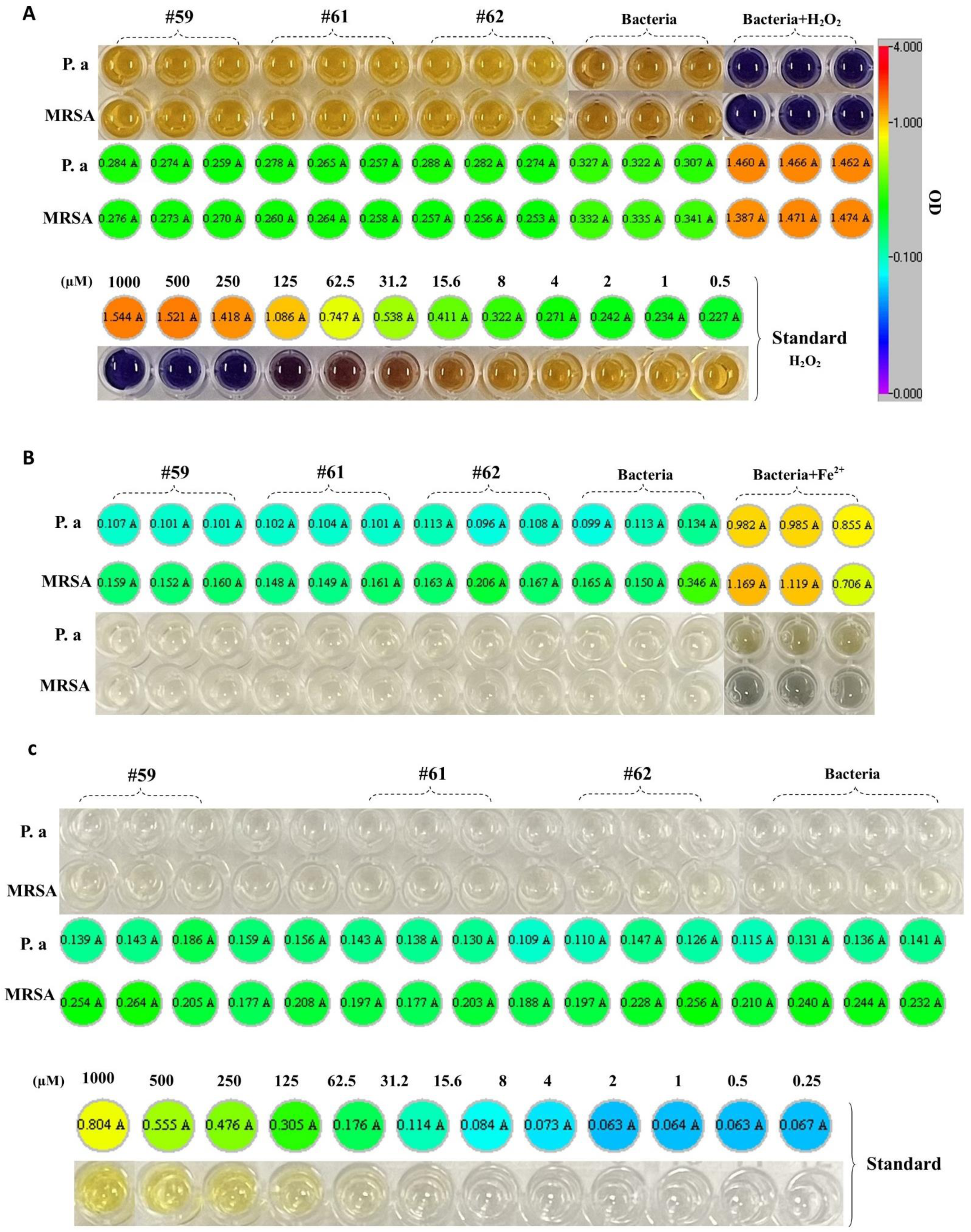
A. Hydrogen peroxide (H_2_O_2_) concentration. Shows the H_2_O_2_ concentrations in the samples with the naked eye (down) and plate reader (upper). The standard, 1mM solution of hydrogen peroxide serially diluted with double distilled (DD) water 1:2 for a total of 11 samples. DD water was used as the blank and working reagent (WR). **B. Free iron [(ferrous) Fe^+2^] concentration.** After 1 h treatment with selected components relative amount of ferrous measured by OD. The naked eye (upper) Heatmap (down) is illustrated in the panels. **C. Reduced thiol (RSH) level.** Shows the RSH absorbance values in the samples with the naked eye (upper) and plate reader (down). 1mM solution of glutathione reduced oxidized serially diluted with Tris/HCl 1:2 for a total of 11 samples as a standard. The 50 mM Tris/HCl pH 8 and Ellman’s reagent were used as the blank.

Due to the considerable structural differences between the Gram-positive and Gram-negative cell membranes of bacteria(18), as well as eukaryotic cells (19), certain antibiotics, such as polycations and chelators, selectively target the distinctive structure of bacterial cell membranes (20). This targeted approach allows antibiotics to eradicate bacteria more specifically while preserving host cells (20). In our investigation of membrane disruption potency within our chosen components, we employed the membrane leakage probe (PI), which binds to DNA, to assess membrane destabilization following a 1-hour exposure to our agents. Results indicate that all selected agents disrupted the Gram-positive cell membrane compared to the untreated group (p < 0.001), but not sig effect of Gram-negative cell membrane.

### Toxicity in Eukaryotic Cell lines

We selected compounds EPB-59, EPB-61, and EPB-62 due to their elevated antibacterial activity. Subsequently, we evaluated their cytotoxicity in eukaryotic cell lines derived from human, canine, or simian origins. In HEL, HeLa, MT4 and VERO cells, the MCC or CC_50_ values were similar for the three compounds and in the order of 4-20 µM. EPB-59 and EPB-61 showed lower cytotoxicity in canine MDCK cells, giving a CC_50_ value ≥100 µM (**Table 3**). The CC50 values of EPB-59 and EPB-61 on MDCK cells reveal their exceptional cytotoxic profiles in comparison to other compounds, with concentrations varying from ≤ 0.8 µM to a > 100 µM. Notably, EPB-59 and EPB-61 possess cytotoxicity levels 2 logs lower than that of EPB-34.

**Table 3.** Eukaryotic cell Toxicity of different diphenyl-urea analogs.

| | Cell line (in vitro cytotoxicity, μM) | | | | |
| --- | --- | --- | --- | --- | --- |
|  | Human |  |  | Mouse | Monkey |
|  | HEL<sup>a</sup><br>(MCC)<sup>d</sup> | HeLa<sup>b</sup><br>(MCC)<sup>d</sup> | MT4<sup>c</sup><br>(MCC)<sup>d</sup> | MDCK<sup>e</sup><br>(CC<sub>50</sub>)<sup>g</sup> | Vero<sup>f</sup><br>(CC<sub>50</sub>)<sup>g</sup> |
| **EBP-59** | >4 | 4 | 3.9 | >100 | >4 |
| **EBP-61** | >4 | 4 | 2.8 | >100 | >4 |
| **EBP-62** | 20 | >4 | 4.7 | 100 | >4 |

<sup>a</sup>HEL human leukemia cells. <sup>b</sup>HeLa human cervical cancer cells. <sup>c</sup>MT4 human T leukemia cells. <sup>d</sup>MCC: minimum compound concentration that causes a microscopically detectable alteration of normal cell morphology. <sup>e</sup>MDCK: Madin-Darby canine kidney cells. <sup>f</sup>Vero: Green monkey kidney cells. <sup>g</sup>50% Cytotoxic concentration, as determined by measuring the cell viability with the colorimetric formazan- based MTS assay. Values shown are the mean of 2 or 3 determinations.

### In vitro results

The new diphenylurea compounds studied here were not effective against Gram-negative pathogens such as *E. coli, P. aeruginosa, K. pneumonia,* and *P. mirabilis* except compound **59**, which had acceptable potency regarding MIC, MBC, and MBIC to *P. mirabilis* spp. This could indicate that variations in diphenyl-urea antibiotics could make them efficient to use against Gram-negative bacteria. Perhaps this compound and how it differs from others could be a future topic for further study. Furthermore, promising results regarding the application of compounds **59**, **61**, and **62** were achieved with Gram-positive bacteria such as *S. aureus, S. epidermidis,* and methicillin-resistant strains *S. aureus* (*MRSA*) specifically with compound **59**, having relatively superior results while factoring cell cytotoxicity in all cell lines evaluated. Even though results regarding the use of such compounds on *MRSA* species did not accompany a safe dose regarding human cell lines, MIC, MBC, and MBIC measurements of 10 *MRSA* clinical isolates showed these compounds could be used in a rather safe measure when evaluated on clinical isolates. All *MRSA* clinical isolates showed susceptibility to compounds **59**, **61** and **62** regarding MBIC and MIC (**Table 2**). With compound **59** yet again proving to be the best option across all measurements regarding safety and efficacy, all these compounds showed a significantly higher potency when compared to conventional antibiotics evaluated.

Other compounds were also evaluated in this study. However, many of them did not prove as effective as the three discussed above. Compound **34** showed promising results regarding MIC in Gram-positive bacteria assessed in this study with a similar cytotoxicity profile as compound **59** regarding HeLa, HEL, and Vero cell lines. Unfortunately, the same does not apply in the context of MBIC and MBC. Compound **37** showed slightly worse results and its cytotoxicity was not assessed. Compound **40** did not show any promising effectiveness against Gram-positive bacteria even in comparison to conventional antibiotics.

### In vivo efficacy

The most effective compounds-**59**, **61**, and **62**, were selected to evaluate their *in vivo* efficacy. Mice were infected systemicallywith the community-associated MRSA *strain* MW2 at an infectious dose of 5×10^7^ CFU as described (21). Treatment groups were injected intraperitoneally with the compound of interest one hour before or after infection, with a saline-injected control group. After 24 hours of infection, mice were sacrificed and blood, peritoneal fluid, spleen, liver, kidney (left), heart, and lungs were harvested for CFU enumeration to determine the bacterial burden in these organs. These experiments revealed a consistently high bacterial burden in all organs of the animals 24 hours after treatment, irrespective of the specific treatment administered (**Fig 4**). The single-dose treatment failed to reduce bacterial counts in any of the animal organs, even after experimenting with different compound concentrations. While disappointing, the lack of promising results in the single-dose treatments may have a variety of reasons including, rapid metabolism and excretion of the compounds by the liver or to the acknowledged intracellular aspect of *S. aureus* in its infectious cycle (22). Consequently, further research is imperative to determine the *in vivo* efficacy of compounds **59**, **61**, and **62**. Specifically, if a pulsatile and continuous treatment over 2-4 days may prove to be more effective. Additionally, investigating the possibility of topical application for these components presents another viable option.

**Fig 4.**
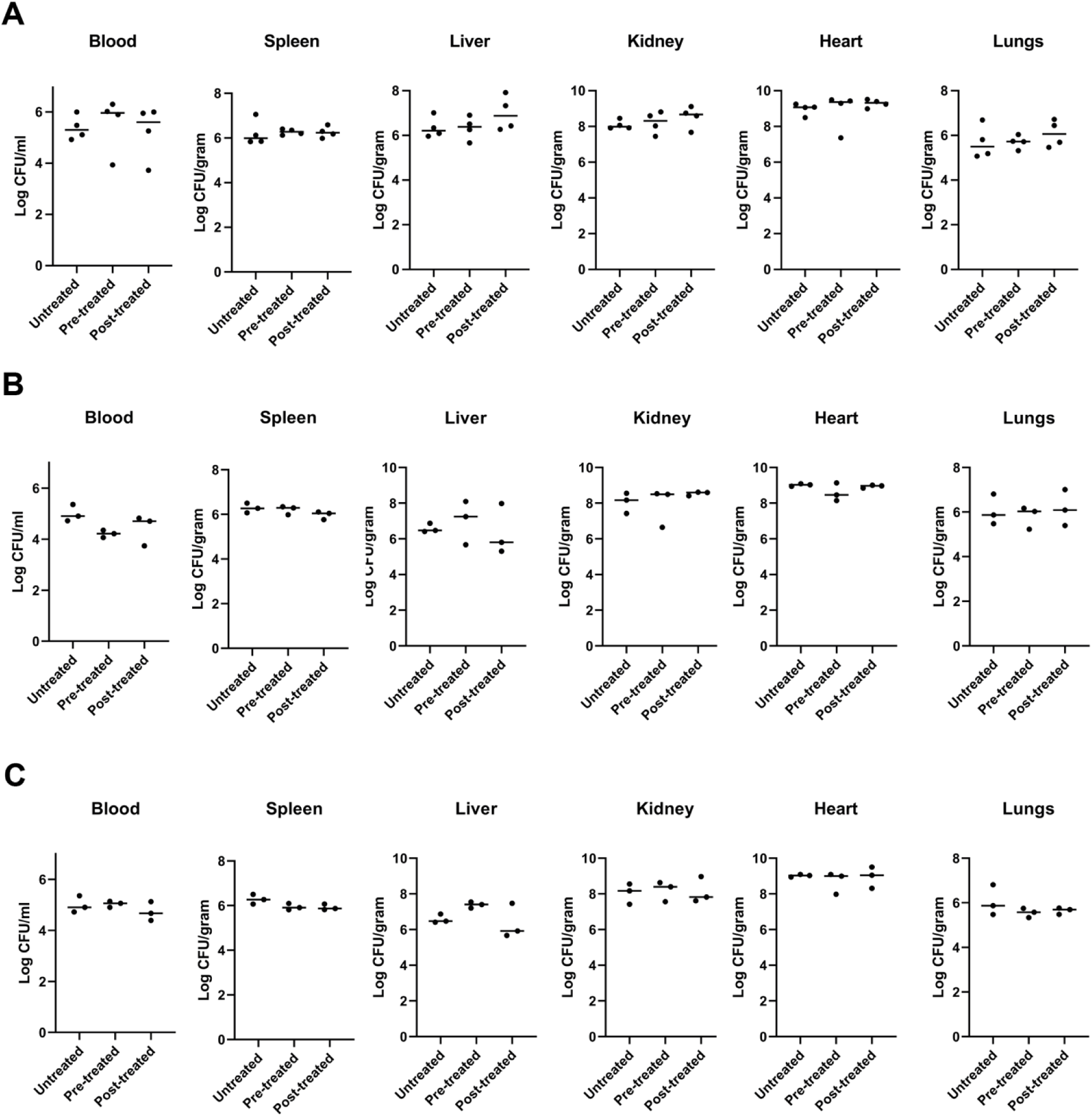
Intravenous MRSA Infection model to screen for *In vivo* effectivity of compound-59, 61, and 62. CFU enumerations for blood, spleen, liver, kidney (left), heart, and lungs after 24 hours of infection with *S. aureus* MW2. Mice were left untreated or were treated 1 hour before infection (pre-treated) or 1 hour after infection (post-treated) with compound **59 (A)**, **61 (B)**, or **62 (C)**. One-way ANOVA with Bonferroni’s posttest, data not significant.

## Conclusion

Overall, for the first evaluation of such compounds, the results demonstrated Gram class selectivity, a cell wall/membrane associated mechanism of antimicrobial activity. Cell toxicity and animal results indicate challenges but still suggest there is promise to explore applications of these novel compounds as useful antimicrobials moving forward.

## Materials and Methods

### Pentafluorosulfanyl-containing triclocarban analogs preparation and characterization

All the pentafluorosulfanyl-containing diarylureas were synthesized as previously reported (5, 12, 13). In all cases the analytical and spectroscopic data matched with those previously reported in the above mentioned three references.

### Bacterial Preparation

All bacterial strains subjected to testing had been stored at −70°C prior to the experiment. The experiment commenced by streaking small amounts of the frozen bacterial samples onto Muller-Hinton agar (MHA) culture medium (BD Bacto, Oxoid, Basingstoke, UK Cat# X296B) and allowing for overnight incubation. Bacteria were re-cultured on MHA plates as required to confirm colony morphology and lack of contamination. Strains tested were *Pseudomonas aeruginosa* ATCC 27853, *Escherichia coli* ATCC 25922, *Staphylococcus aureus* ATCC 25923, Methicillin resistant *S. aureus* (*MRSA*) ATCC 33591, *Staphylococcus epidermidis* ATCC 12228, *Klebsiella pneumoniae* ATCC, *Proteus mirabilis* ATCC 35659 and 10 clinical isolates of *MRSA* from Foothills hospital, Calgary, Canada. To prepare for susceptibility testing, a protocol modified from Lemire *et al*was used (23). MHB was swab inoculated from MHA colonies and shake incubated until turbid. Dilution of the broth with 0.9% saline was performed as necessary to match a 1.0 McFarland standard.

### Planktonic Susceptibility

Minimum inhibitory concentration (MIC) and minimum bactericidal concentration (MBC) assessments were conducted on all bacterial strains using 96-well microtiter plates. To prepare the wells for bacteria, serial dilutions of the antibacterial agents were carried out along the rows of two microtiter plates. The antibacterials were 2-fold diluted with Mueller-Hinton broth (MHB) in each column, reaching a volume of 75 µL. The 1.0 McFarland standard was diluted 15-fold in MHB, and 75 µL of inoculum was added to the wells treated with antibacterials, resulting in a total volume of 150 µL. The microtiter plates were covered, then shake-incubated overnight at 150 rpm and 37°C. MIC was determined based on visible bacterial growth. In cases of antibacterial opacity or ambiguous results, streaking of well cultures on Mueller-Hinton agar (MHA), overnight incubation, and subsequent comparison to controls were performed. MBC was determined by transferring 3 µL of culture from all MIC microtiter plate wells to 147 µL of Mueller-Hinton broth in fresh plates. These plates were shake-incubated overnight at 150 rpm and 37°C, and bacterial growth was visually inspected.

### Inhibition of Biofilm formation

Minimum biofilm inhibitory concentration (MBIC) determination coincided with MIC and MBC assessments. The Calgary Biofilm Device (CBD) was employed to cover the 150 µL wells of the MIC microtiter plate. Following overnight incubation, the MIC plates and pegs underwent three washes with 250 µL of distilled water (dH_2_O) to eliminate planktonic bacteria. Biofilms were then stained with 200 μL of a 0.1% crystal violet solution for 30 minutes. Post-staining, microplates and pegs were washed three times with 200 μL of ddH_2_O to remove excess dye. Biofilm quantification involved sonication using a 250HT ultrasonic cleaner (VWR International), set at 60 Hz for 10 minutes, into 200 μL of 70% ethanol. Absorbance readings at 600 nm were taken, with 70% ethanol serving as the blank (1).

### Membrane Permeability Measurements

For assessing membrane permeability, propidium iodide (PI) (Invitrogen, Eugene, Oregon, USA) served as the fluorescent reporter dye. Increased PI fluorescence readings are indicative of heightened membrane disruption and permeability, as PI can penetrate cells, bind to DNA, and remain within the cells (24, 25). Bacteria were cultured in 3 mL of Mueller-Hinton broth (MHB) and incubated at 37 °C for approximately 3 hours in a shaker incubator (150 rpm) until reaching an OD600 of 0.08. Each group received treatment with MIC concentrations of agents, the untreated groups with phosphate-buffered saline (PBS) as a negative control, and a positive control involving bacteria subjected to freezing and boiling three times (frozen for 5 minutes at −80 °C and incubated at 90 °C for 10 minutes) to disrupt the bacterial cell membrane. All treated and control groups were incubated at 37 °C for 1 hour in a shaker incubator (150 rpm).

Ciprofloxacin and gentamycin were employed as internal controls, as they are known not to target the bacterial cell membrane in the initial steps. After incubation, the samples were centrifuged, washed with PBS (10,000 rpm for 2 minutes), and bacteria were stained with 0.08 mM PI for 5 minutes at 21 °C in the dark. Subsequently, 10 μL of samples were transferred onto slides and examined under a fluorescence microscope (Zeiss Axio Imager Z1) using the same exposure time (640 ms red, 1s green). Densitometry analysis was conducted using Fiji software (ImageJ).

### Hydrogen Peroxide Assay

The hydrogen peroxide concentration after exposure with our agents was detected with Pierce Quantitative Peroxide Assay Kit, in the aqueous-compatible formulation according to the manufacturer’s instructions (26). For preparing the standard, 1mM solution of H_2_O_2_ was initially made by diluting a 30% H_2_O_2_ stock 1:9000 (11μL of 30% H_2_O_2_ into 100mL of double-distilled (DD) water. This sample was then serially diluted with DD water 1:2 (100μL of DD water + 100μL of the previous dilution) for a total of 11 samples as a standard. 200μL of the working reagent (WR) from the kit was added to 20μL of the diluted H_2_O_2_ standards. Samples were mixed and incubated for 15 minutes at 21 °C in the dark. Absorbances were measured at 595nm using a Thermomax microtiter plate reader with Softmax Pro data analysis software (Molecular Devices, Sunnyvale, CA).

For measuring the treated and untreated samples. The bacteria were cultured in 2mL of MHB and were incubated at 37 °C for ∼ 3 h in a shaker incubator (150 rpm) to reach the OD600 of 0.08. Then treated with MIC concentrations of agents and untreated groups with PBS as a negative control, the positive control was treated with 250μM H_2_O_2,_ and incubated at 37 °C for 1 h in a shaker incubator (150 rpm). Ciprofloxacin and gentamycin were used as antibiotic comparators. The bacterial cells were washed with PBS by centrifuging (10,000 rpm for 5 min) and discarding the supernatant. 2mL PBS was added to each sample and vortexed. 200μL of the WR was added to 20μL of each sample. Samples were mixed and incubated for 15 minutes at room temperature. Absorbances were measured at 595nm using a Thermomax microtiter plate reader with Softmax Pro data analysis software (Molecular Devices, Sunnyvale, CA). The blank value was subtracted from all sample measurements. The samples’ H_2_O_2_ concentrations were calculated based on standard carve R^2^= 0.93 value.

### Iron Detection Ferene-S Assay

The release of Fe^2+^ from the iron-sulfur clusters in *P. aeruginosa* and *MRSA* was detected using a Ferene-S assay with the probe, 3-(2pyridyl)-5,6-bis(2-(5-furylsulfonic acid))-1,2,4-triazin (Sigma-Aldrich, St Louis, MO, USA) (27). The 10 mL of bacteria (OD600 of 0.08) were prepared in Tris-HCl buffer (20 mM, pH 7). The bacterial cells were washed with the same buffer by centrifuging (10,000 rpm for 5 min) and discarding the supernatant. The platelet (bacterial cells) was then lysed by sonication using a 250HT ultrasonic cleaner (VWR International), set at 60 Hz for 20 min in the same buffer. The samples were centrifuged (10,000 rpm for 5 min) and the supernatant was collected. The solution was treated with MIC concertation of agents, untreated control (PBS) and positive control (incubated in 90 °C for 10 min to break down the Fe-sulfur cluster). Then, a 10 mM Ferene-S probe was added to each sample in a 96 well plate, and samples were incubated at 21 °C in dark for 1 h. Absorbance was measured at 600 nm, using a Thermomax microtiter plate reader with Softmax Pro data analysis software (Molecular Devices, Sunnyvale, CA) (24).

### Reduced thiol (RSH) assay

The accuracy of the assay was assessed with a standard dilution of reduced glutathione (≥98%, Alfa Aesar, Germany) and oxidized glutathione (Sigma, USA). For preparing the standard, 1mM solution of each glutathione was serially diluted with 50 mM Tris/HCl pH 81:2 (150μL of 50 mM Tris/HCl pH 8 + 150μL of glutathione) for a total of 11 samples as a standard. Then 0.1 mM of Ellman’s reagent 5,5’-dithiobis(2-nitrobenzoic acid) (DTNB) was added to each well. Samples were mixed and incubated for 30 minutes at 37 °C in dark. Absorbances were measured at 412nm using a Thermomax microtiter plate reader with Softmax Pro data analysis software (Molecular Devices, Sunnyvale, CA).

For measuring the treated and untreated samples with agents. The bacteria were cultured in 3mL of MHB and were incubated at 37 °C for ∼ 3 h in a shaker incubator (150 rpm) to reach the OD600 of 0.08. Then treated with MIC concentrations of agents and untreated groups with PBS (as a negative control) and incubated at 37 °C for 1 h in a shaker incubator (150 rpm). The bacterial cells were washed with PBS by centrifuging (10,000 rpm for 5 min) and discarding the supernatant. 1mL of 50 mM Tris/HCl pH 8.0, 5 mM EDTA, 0.1% SDS and 0.1 mM DTNB was added to each sample and vortexed. These cell suspensions were incubated at 37°C for 30 min, and then centrifuged in a microfuge for 1 min at 15000 g. Absorbances were measured at 412nm using a Thermomax microtiter plate reader with Softmax Pro data analysis software (Molecular Devices, Sunnyvale, CA). The blank value was subtracted from all sample measurements. The absorption coefficient of oxidized DTNB (1.36¬10^4^ M^-1^ cm^-1^) at this wavelength was then used to calculate the RSH concentration of the cell.

### Eukaryotic cell toxicity of pentafluorosulfanyl-containing triclocarban compounds

Five Eukaryotic cell lines were used to determine the cytotoxicity of the compounds: human embryonic lung (HEL) fibroblast cells; human cervixcarinoma-derived HeLa cells; human T-cell leukemia-derived MT4 cells; Madin-Darby canine kidney (MDCK) cells; and African Green monkey kidney-derived VERO cells. Semi-confluent cell cultures in 96-well plates were exposed to serial dilutions of the compounds or to medium (= no compound control), then incubated at 37 °C. Four days later, the cells were inspected by microscopy to determine the Minimum Cytotoxic Concentration (MCC), i.e. compound concentration that causes a microscopically detectable alteration of normal cell morphology. Next, the MTS cell viability reagent (CellTiter 96 AQueous MTS Reagent from Promega) was added. After 4 h incubation at 37 °C, optical density (OD) at 490 nm was recorded in a microplate reader. The percentage cytotoxicity was calculated as: [1 - (OD_Cpd_)/(OD_Contr_)] x 100, after which the 50% cytotoxicity value (CC_50_) was derived by extrapolation, assuming semi-log dose response.

### Bacterial infection and mouse treatment

Animal experiments were performed with 6-10 week/old adult male mice. Mice were kept in a pathogen-free facility under standardized conditions: temperature of 21-22 °C, illumination of 12h light-12h dark, and access to tap water and food. Experimental animal protocols were approved by the University of Calgary Animal Care Committee and followed the Canadian Council for Animal Care Guidelines.

MW2 *Staphylococcus aureus* were grown on brain heart infusion (BHI) agar plates (BD biosciences). A single colony was grown overnight in BHI medium (BD biosciences) at 37 °C while shaking. Subcultures of 100 ml of the ON culture were grown in 3 ml of the same culture medium for 2h at 37 °C while shaking to reach the exponential growth phase. Bacteria were brought to a concentration of OD_660_ 1.0 in saline. A 1:4 dilution of bacteria was prepared, and animals were infected through an intravenous catheter with 200 ml of the bacterial suspension, so that bacterial dose was controlled to a dose of 5×10^7^ bacteria. Mice were treated with and intraperitoneal injection of 100 µl of compound of interest (5 mM) 1 hour before infection or 1 hour after infection. Untreated group was injected 1-hour post-infection with 100 µl saline.

For assessing bacterial by CFU enumeration, the mice were anesthetized with isoflurane (Fresenius Kabi) and washed with 70% ethanol to collect blood, and sacrificed to further collect liver, spleen, kidney (left), heart, and lungs. Blood was collected by cardiac puncture and pooled with 40 ml heparin. 25 ml of 1:10 diluted blood samples were plated. Organ samples were weighed and homogenized in 1 ml PBS and serially diluted (spleen, liver, and lung samples 1:10, 1:100; 1:1000; 1:10 000; kidney and heart samples 1:100; 1:1000; 1:10 000; 1:100 000). 30 ml of each dilution was plated on a 30° angle to form stripes. All plates were incubated ON at 37 °C. Cultures were counted, and colony-forming units (CFUs) were calculated taking the weight of the organ into account.

## Acknowledgements

RJT supported by a Discovery Grant from the Natural Sciences Engineering Research Council of Canada. BGJS is supported by the Canada Research Chair (tier2) program.

Grant PID2021-122116OB-I00 (M.V-C.) funded by MCIN/AEI/10.13039/501100011033 and by “ERDF A way of making Europe”. CIBER de Diabetes y Enfermedades Metabólicas Asociadas (CIBERDEM) is a Carlos III Health Institute project. CERCA Programme/Generalitat de Catalunya.

## Author contributions

Conceived and designed the study: **AP, MVC, RJT**

Practical performance: **AP, MM, JWH, EP, LN, SV, BGS, MZ, MVC, RJT**

Analyzed the data: **AP, MM, JWH, MVC, RJT**

Wrote the paper: **AP, MM, MVC, RJT**

Participated in data analysis and manuscript editing: **AP, MM, JWH, EP, LN, SV, BGS, MZ, MVC, RJT**

All authors **AP, MM, JWH, EP, LN, SV, BGS, MZ, MVC, RJT** have read and agreed to the published version of the manuscript.

## References

1. Getahun H, Smith I, Trivedi K, Paulin S, Balkhy HH. 2020. Tackling antimicrobial resistance in the COVID-19 pandemic. Bulletin of the World Health Organization 98:442.

2. Pushpakom S, Iorio F, Eyers PA, Escott KJ, Hopper S, Wells A, Doig A, Guilliams T, Latimer J, McNamee C. 2019. Drug repurposing: progress, challenges and recommendations. Nature reviews Drug discovery 18:41–58.

3. Catalano A, Iacopetta D, Pellegrino M, Aquaro S, Franchini C, Sinicropi MS. 2021. Diarylureas: Repositioning from Antitumor to Antimicrobials or Multi-Target Agents against New Pandemics. Antibiotics 10:92.

4. Farha MA, Brown ED. 2019. Drug repurposing for antimicrobial discovery. Nature microbiology 4:565–577.

5. Pujol E, Blanco-Cabra N, Julián E, Leiva R, Torrents E, Vázquez S. 2018. Pentafluorosulfanyl-containing triclocarban analogs with potent antimicrobial activity. Molecules 23:2853.

6. Altomonte S, Zanda M. 2012. Synthetic chemistry and biological activity of pentafluorosulphanyl (SF5) organic molecules. Journal of Fluorine Chemistry 143:57–93.

7. Savoie PR, Welch JT. 2015. Preparation and utility of organic pentafluorosulfanyl-containing compounds. Chemical reviews 115:1130–1190.

8. Bassetto M, Ferla S, Pertusati F. 2015. Polyfluorinated groups in medicinal chemistry. Future Medicinal Chemistry 7:527–546.

9. Gujjar R, El Mazouni F, White KL, White J, Creason S, Shackleford DM, Deng X, Charman WN, Bathurst I, Burrows J, Floyd DM, Matthews D, Buckner FS, Charman SA, Phillips MA, Rathod PK. 2011. Lead optimization of aryl and aralkyl amine-based triazolopyrimidine inhibitors of Plasmodium falciparum dihydroorotate dehydrogenase with antimalarial activity in mice. J Med Chem 54:3935–49.

10. Moraski GC, Bristol R, Seeger N, Boshoff HI, Tsang PS, Miller MJ. 2017. Preparation and Evaluation of Potent Pentafluorosulfanyl-Substituted Anti-Tuberculosis Compounds. ChemMedChem 12:1108–1115.

11. Morehead MS, Scarbrough C. 2018. Emergence of global antibiotic resistance. Primary care: clinics in office practice 45:467–484.

12. Probst A, Pujol E, Häberli C, Keiser J, Vázquez S. 2021. In vitro, in vivo, and absorption, distribution, metabolism, and excretion evaluation of SF5-Containing N, N’-diarylureas as antischistosomal agents. Antimicrobial agents and chemotherapy 65:10.1128/aac.00615-21.

13. Zarei M, Pujol E, Quesada-López T, Villarroya F, Barroso E, Vázquez S, Pizarro-Delgado J, Palomer X, Vázquez-Carrera M. 2019. Oral administration of a new HRI activator as a new strategy to improve high-fat-diet-induced glucose intolerance, hepatic steatosis, and hypertriglyceridaemia through FGF21. British journal of pharmacology 176:2292–2305.

14. Wu H, Moser C, Wang HZ, Høiby N, Song ZJ. 2015. Strategies for combating bacterial biofilm infections. Int J Oral Sci 7:1–7.

15. Ceri H, Olson ME, Stremick C, Read RR, Morck D, Buret A. 1999. The Calgary Biofilm Device: New Technology for Rapid Determination of Antibiotic Susceptibilities of Bacterial Biofilms. Journal of Clinical Microbiology 37:1771–1776.

16. Reimche JL, Kirse DJ, Whigham AS, Swords WE. 2016. Resistance of non-typeable Haemophilus influenzae biofilms is independent of biofilm size. Pathogens and Disease 75.

17. Wannigama DL, Hurst C, Hongsing P, Pearson L, Saethang T, Chantaravisoot N, Singkham-In U, Luk-In S, Storer RJ, Chatsuwan T. 2020. A rapid and simple method for routine determination of antibiotic sensitivity to biofilm populations of Pseudomonas aeruginosa. Annals of Clinical Microbiology and Antimicrobials 19:1–8.

18. de CC Pinto N, Campos LM, Evangelista ACS, Lemos AS, Silva TP, Melo RC, de Lourenco CC, Salvador MJ, Apolonio ACM, Scio E. 2017. Antimicrobial Annona muricata L.(soursop) extract targets the cell membranes of Gram-positive and Gram-negative bacteria. Industrial Crops and Products 107:332–340.

19. Theriot JA. 2013. Why are bacteria different from eukaryotes? BMC biology 11:1–17.

20. Vaara M. 1992. Agents that increase the permeability of the outer membrane. Microbiological reviews 56:395–411.

21. Surewaard BG, Deniset JF, Zemp FJ, Amrein M, Otto M, Conly J, Omri A, Yates RM, Kubes P. 2016. Identification and treatment of the Staphylococcus aureus reservoir in vivo. Journal of Experimental Medicine 213:1141–1151.

22. Hommes JW, Surewaard BG. 2022. Intracellular habitation of Staphylococcus aureus: Molecular mechanisms and prospects for antimicrobial therapy. Biomedicines 10:1804.

23. Lemire JA, Kalan L, Gugala N, Bradu A, Turner RJ. 2017. Silver oxynitrate–an efficacious compound for the prevention and eradication of dual-species biofilms. Biofouling 33:460–469.

24. Morones-Ramirez JR, Winkler JA, Spina CS, Collins JJ. 2013. Silver enhances antibiotic activity against gram-negative bacteria. Science translational medicine 5:190ra81–190ra81.

25. Novo DJ, Perlmutter NG, Hunt RH, Shapiro HM. 2000. Multiparameter flow cytometric analysis of antibiotic effects on membrane potential, membrane permeability, and bacterial counts of Staphylococcus aureus and Micrococcus luteus. Antimicrobial agents and chemotherapy 44:827–834.

26. https://www.thermofisher.com/document-connect/document-connect.html?url=https%3A%2F%2Fassets.thermofisher.com%2FTFS-Assets%2FLSG%2Fmanuals%2FMAN0011275_Pierce_Quant_Peroxide_Asy_UG.pdf.

27. Hennessy DJ, Reid GR, Smith FE, Thompson SL. 1984. Ferene—a new spectrophotometric reagent for iron. Canadian journal of chemistry 62:721–724.

28. Zarei M, Pujol E, Quesada-López T, Villarroya F, Barroso E, Vázquez S, Pizarro-Delgado J, Palomer X, Vázquez-Carrera M. 2019. Oral administration of a new HRI activator as a new strategy to improve high-fat-diet-induced glucose intolerance, hepatic steatosis, and hypertriglyceridaemia through FGF21. Br J Pharmacol 176:2292–2305.

